# Hyperglycemia-Driven Hepatic Immune Dysfunction Facilitates Microbial Dissemination Post-Myocardial Infarction

**DOI:** 10.1101/2025.02.26.640249

**Authors:** Tony W.H. Tang, Sumi Nani Novita Pasaribu, Hung-Chih Chen, Po-Ju Lin, Shu-Chian Ruan, Emily Wissel, Chiung-Wen Kuo, Peilin Chen, Yen-Chun Ho, Yi-Cheng Chang, Patrick C.H. Hsieh

## Abstract

**Background:** The gut microbiome is intimately connected to cardiovascular health through the gut-heart axis and plays a pivotal role in maintaining homeostasis. Myocardial infarction (MI) disrupts this homeostatic balance, leading to widespread adverse effects. Hyperglycemia, a hallmark of metabolic dysfunction, further exacerbates these disruptions, emphasizing the need to understand the underlying mechanisms to develop effective therapeutic strategies for mitigating the cascading complications along the gut-heart axis. This study aims to elucidate the dynamics of gut barrier disruption during MI, and explore the liver’s function as an immune sentinel in this process, with a focus on the impact of hyperglycemia on microbial dissemination, systemic inflammation, and liver immune function.

**Methods:** A murine MI model was used to evaluate gut permeability, bacterial translocation, and hepatic immune responses. MI was induced via permanent left anterior descending artery ligation. Hyperglycemia was established through streptozotocin injections and a high-fat, high-sugar diet. Gut barrier integrity was assessed using FITC-dextran assays, and microbial translocation was tracked through intravital imaging and anaerobic bacterial cultures from multiple organs. Hepatic immune function was analyzed via flow cytometry, cytokine profiling, and phagocytosis assays. 16S rRNA sequencing characterized the composition of translocated bacteria.

**Results:** MI significantly increased intestinal permeability, with hyperglycemia further exacerbating gut barrier dysfunction. Intravital imaging revealed bacterial translocation through the portal vein to the liver, highlighting the liver’s role in microbial interception. Hyperglycemia impaired hepatic macrophage function by activating NLRP3 inflammasome signaling, reducing bacterial clearance and promoting persistent liver colonization. Systemic inflammatory cytokines, particularly TNF-α, were elevated, further facilitating microbial dissemination. 16S rRNA sequencing demonstrated host-dependent stochastic variability in translocated bacterial composition.

**Conclusion:** The liver serves as a key immune regulator in the gut-liver-heart axis but is functionally compromised under hyperglycemia, exacerbating systemic inflammation and microbial dissemination post-MI. Targeting NLRP3 signaling and restoring gut barrier integrity may mitigate post-MI complications, particularly in hyperglycemic conditions. These findings underscore the need for integrated therapeutic strategies incorporating metabolic control and microbiome-targeted interventions to improve post-MI outcomes.

## Introduction

The human microbiome, an intricate ecosystem comprising trillions of microorganisms, plays a crucial role in regulating host physiology and maintaining systemic homeostasis [1,2]. This symbiotic relationship has been integral to human evolution and profoundly influences health and disease states across various organ systems [3–8].

Cardiovascular diseases (CVD), particularly myocardial infarction (MI), pose a significant global health burden [9]. MI is characterized by ischemia-induced myocardial necrosis, often leading to systemic complications beyond the cardiac tissue [10,11]. Emerging evidence highlights the interconnectedness of the gut microbiome and cardiovascular health, commonly referred to as the gut-heart axis [12–16]. Recent studies have underscored the pivotal role of the gut microbiome in cardiac health and its disruption in the context of MI [17,18]. Disruptions in the gut microbial composition and integrity have been implicated in increased risk post-MI and poorer clinical outcomes, with experimental models demonstrating elevated intestinal permeability following cardiac injury [19,20]. Loss of gut microbiota homeostasis after MI exacerbates intestinal permeability, leading to the systemic spread of bacterial components that drive inflammation and impair recovery after injury [18].

Furthermore, the depletion of gut-derived short-chain fatty acids (SCFAs), such as butyrate, has been linked to increased systemic inflammation and hindered cardiac repair post-MI [14]. Similarly, the compromised gut barrier also enables the translocation of gut-derived bacteria and inflammatory mediators, such as lipopolysaccharides (LPS), into systemic circulation [19]. This microbial leakage exacerbates systemic inflammation, amplifies cardiac dysfunction, and hinders recovery and survival outcomes.

Notably, facultative anaerobic gut bacteria have been shown to colonize hypoxic regions such as solid tumors, following systemic dissemination [21–23]. The hypoxic and necrotic myocardial environment resulting from MI provides a similar niche for these bacteria [24,25], suitable for their colonization in cardiac tissue after systemic leakage. However, whether specific bacterial species preferentially colonize post-MI tissue, thereby contributes to exacerbated cardiac inflammation and impaired repair, remains to be explored.

Hyperglycemia, which has emerged as a critical factor in shaping post-MI outcomes, further complicates the interplay between disrupted health states and microbial states [26,27]. Persistent hyperglycemia is strongly associated with increased mortality and complications, with elevated glucose levels following Acute MI (AMI) significantly affecting survival rates [28]. Moreover, intensive glucose control in diabetic patients post-AMI has been shown to reduce mortality, reinfarction, and heart failure [29,30]. Beyond persistent hyperglycemia, glycemic variability—characterized by fluctuations in blood glucose levels—has been identified as an independent predictor of adverse cardiovascular outcomes [31]. Even transient hyperglycemia during the acute phase of AMI in non-diabetic individuals is linked to higher mortality and complication rates [31]. Furthermore, hyperglycemia escalates gut permeability and diminishes immune defenses in other organs, such as the liver, particularly the macrophage-mediated interception of disseminated microbes [32–35]. The cross-organ dysregulation caused by systemic metabolic conditions profoundly influences outcomes following cardiac injury. These findings highlight the systemic impact of glucose dysregulation and underscore the necessity of comprehensive management strategies to optimize post-MI recovery and outcomes.

Our study delves into the complex processes underlying microbial dynamics and cross-organs interactions in the aftermath of MI. Specifically, it investigates how hyperglycemia, a common feature of metabolic dysfunction, may modulate microbial behavior and contribute to systemic inflammation. Understanding these mechanisms is crucial for developing targeted therapeutic strategies to address the interconnected inflammatory and microbial challenges associated with cardiac injury and metabolic dysfunction. This research is essential for advancing our understanding of how systemic conditions and cross-organs interactions influence cardiac health and how they can be effectively managed.

## Materials and Methods

### Animals

C57BL/6 specific pathogen-free (SPF) mice were obtained from the National Laboratory Animal Center (NLAC), Taiwan. The mice were housed under controlled conditions with a 12-hour light/dark cycle and provided ad libitum access to sterile chow diet (Cat No. 5053; LabDiet, USA) and water. All experimental procedures involving mice were approved by the Institutional Animal Care and Use Committees (IACUC) of both Academia Sinica and NLAC (Approval No. 24022137). To ensure unbiased results, the surgeon conducting the procedures was blinded to the experimental groups, and mice were randomly assigned to the respective groups prior to surgery.

### Surgery and Heart Function Assessment

Myocardial infarction (MI) was induced by permanent ligation of the left anterior descending coronary artery, performed 2–3 mm distal to the left atrial appendage, as previously described [18]. Cardiac function was assessed three days post-MI using echocardiography, conducted with a Vivid-q Ultrasound system (GE Healthcare, USA) equipped with a 5–13 MHz intraoperative probe.

### Antibiotic Treatment

To induce dysbiosis in SPF mice, a broad-spectrum antibiotic cocktail was prepared by dissolving ampicillin (0.25 g/L), metronidazole (0.25 g/L), neomycin (0.25 g/L), and vancomycin (0.125 g/L) (all from Sigma-Aldrich, USA) in autoclaved water. The cocktail was administered to the mice starting seven days prior to surgery to ensure the establishment of gut microbial disruption.

### Development of STZ-induced Hyperglycemia Model

Mice were intraperitoneally injected with streptozotocin (STZ) (Cat. S0130, Sigma-Aldrich, USA) (50 mg/kg body weight) daily for five consecutive days. Following the injections, the mice were fed a high-fat, high-sugar diet (HFHSD) (Cat. D12331, Research Diets, USA) for one week. Blood samples were collected from the tail vein and tested using a portable glucose meter (Roche, USA). Mice with blood glucose levels exceeding 300 mg/dL were classified as hyperglycemic and included in the study.

### Epicardium Injection and Colonization of Stool-isolated Bacteria

A stool bacterial culture was prepared in Brain Heart Infusion (BHI) (Cat. 221812, BD, USA) broth and incubated overnight at 37°C under anaerobic conditions. Bacteria were harvested by centrifugation at 5,000 × g for 10 minutes, and the pellet was re-suspended in sterile phosphate-buffered saline (PBS) to achieve the desired concentration. A left thoracotomy was performed on mice to expose the heart. Using a 31G insulin syringe, 10 μl of the prepared bacterial suspension was injected directly onto the epicardium. The chest wall was then closed with absorbable sutures and the skin with non-absorbable sutures. Four days after injection, the mice were euthanized, and their hearts were harvested. The tissues were cultured in sterile tubes containing 3 mL of sterile BHI broth. The tubes were incubated at 37°C under anaerobic conditions, and turbidity was monitored as an indicator of microbial growth. Sterile BHI broth was used as a negative control to confirm the absence of contamination during preparation and handling.

### Detection of Bacteria in Multiple Mouse Organs Using the Broth Culture Method

Organs (e.g., liver, spleen, heart, lungs) were harvested aseptically from euthanized mice and immediately placed in sterile containers on ice to minimize contamination. Each organ was then transferred to a sterile tube containing 3 mL of sterile BHI broth. The tubes were incubated at 37°C under anaerobic conditions, and cultures were monitored for turbidity as an indicator of microbial growth. Sterile BHI broth was included as a negative control to confirm the absence of contamination during preparation and handling.

### Protocol for FITC-dextran-based Gut Permeability Assay

Mice were fasted for 4–6 hours with water provided ad libitum during the fasting period. A FITC-dextran solution (Cat. SI-FD4, Sigma-Aldrich, USA) was prepared at a concentration of 100 mg/mL in sterile PBS and administered orally via gavage. Four hours following FITC-dextran administration, blood samples were collected from the tail vein into heparinized tubes to prevent coagulation. The blood was centrifuged at 2,000 × g for 10 minutes at 4°C, and plasma was collected. The fluorescence intensity of the plasma samples was measured using a microplate reader set to an excitation wavelength of 485 nm and an emission wavelength of 530 nm. A standard curve was generated by performing serial dilutions of the FITC-dextran solution (e.g., 0.1–10 μg/mL in PBS) and was used to quantify the FITC-dextran concentration in the plasma samples.

### Stool DNA Extraction

DNA was extracted from frozen fecal samples using the bead-beating method. Stool DNA from mouse samples was isolated with the Easy-Prep Stool Genomic DNA Kit (Cat No. DPT-BC28; Biotools, Taiwan), following the manufacturer’s protocol. After the final wash step, DNA was eluted in TE buffer and stored at −80 °C for subsequent analysis.

### 16s Ribosomal DNA Sequencing

Full-length 16S rRNA genes were amplified by PCR using barcoded primers (forward: 5’-Phos/GCATC-[16-base barcode]-AGRGTTYGATYMTGGCTCAG-3’ and reverse: 5’-Phos/GCATC-[16-base barcode]-RGYTACCTTGTTACGACTT-3’; degenerate bases: R = A, G; Y = C, T; M = A, C) and KAPA HiFi HotStart ReadyMix. PCR products were analyzed by electrophoresis on a 1% agarose gel, and those with a prominent band at ∼1500 bp were purified using AMPure PB Beads for SMRTbell library preparation. Pooled amplicon libraries were sequenced in circular consensus sequence (CCS) mode on a PacBio Sequel IIe instrument to generate HiFi reads with a Phred quality score ≥30. CCS reads were processed with SMRT Link software, followed by divisive amplicon denoising algorithm 2 (DADA2) (v1.10.1) to obtain amplicons with single-nucleotide resolution from full-length 16S rRNA genes [36]. Amplicon sequences were annotated for taxonomic classification using the GTDB database (r207) via QIIME2 (v2020.11.1) [37]. Further analysis of taxonomic diversity and differential abundance was conducted using the Galaxy platform [34] and MicrobiomeAnalyst [38], incorporating LEfSe (Linear Discriminant Analysis Effect Size) for biomarker identification.

### Histology Examination

Paraffin-embedded tissue sections were processed through a series of steps, including dewaxing and rehydration, followed by staining protocol as briefly described here. To assess intestinal pathophysiology, rehydrated gut sections were incubated with mouse anti-claudin-5 antibodies (Cat No. 34-1600, ThermoFisher, USA). The signal was detected using horseradish peroxidase (HRP)-conjugated anti-mouse IgG antibodies (Cat No. SA00001-7L, Proteintech, USA). For Alcian Blue staining, sections were incubated with 1% Alcian Blue 8GX solution prepared in 0.1 N HCl (pH 1.0) for 2 hours at room temperature. After thorough rinsing in running tap water to remove excess stain, the sections were counterstained with Nuclear Fast Red solution for 1–2 minutes and rinsed in distilled water. Images were captured using an auto scanner, Pannoramic 250 FLASH II (Carl Zeiss, Germany). Quantitative analysis was performed with ImageJ software, ensuring consistent adjustments to brightness and contrast across all images within a series to enhance visual clarity.

### Intravital Imaging

One day post-MI, C57BL/6 mice were intravenously injected with 100 μL of 1 mM FITC-D-alanine (Cat No. SBR00049, Sigma-Aldrich, USA) via the tail vein to label circulating bacteria (n=3). Two hours post-injection, the mice were anesthetized with 2.5% isoflurane, and a laparotomy was performed to expose the portal vein. Images were acquired using an FVMPE-RS multimode multiphoton scanning microscope (Olympus).

### Cytokine Array Assay

Systemic inflammatory cytokine levels were measured using the ProcartaPlex™ Multiplex Immunoassay (Invitrogen, USA), following the manufacturer’s instructions. Plasma samples were collected on day 4 post-MI. The assay was conducted as per the protocol provided and analyzed using the Luminex 100/200 system.

### Clodronate Liposome-mediated Depletion of Hepatic Mononuclear Cells

To deplete hepatic mononuclear cells, mice were intravenously injected with 200 μl of clodronate liposome suspension (Cat. C-010, Liposoma, Netherlands) via the lateral tail vein. Mice were sacrificed two days post-injection for further analysis.

### Isolation of Hepatic Non-parenchymal Cells

Livers were perfused with 5 mL of Hank’s balanced salt solution (HBSS), followed by 12 mL of digestion buffer (HBSS supplemented with 0.5 mg/mL type IV collagenase [Cat. 17104019, Sigma-Aldrich, USA] and 10 μg/mL DNase I [Cat. 10104159001, Roche, USA]). The livers were then excised, minced, and incubated in the digestion buffer at 37°C for 45 minutes. Following digestion, the tissue was passed through a 40 μm cell strainer. Parenchymal cells were removed by low-speed centrifugation at 50 × g for 2 minutes, and the resulting supernatant was enriched with non-parenchymal cells.

### Flow Cytometric Analysis of Immune Cell Composition

Samples were stained with fluorochrome-conjugated antibodies targeting surface or intracellular markers to identify specific cell populations. Surface staining was performed in 2% FBS/PBS for 30 minutes at room temperature. Flow cytometry analysis was conducted using an LSR II-17 flow cytometer. The antibodies used were CD45-FITC, F4/80-APCcy7, and CD11b-AF700 (all from eBioscience, USA). Hepatic mononuclear cells were identified as CD45⁺CD11b⁺F4/80⁺.

### Primary Macrophage Isolation

Mice were intraperitoneally injected with 1 mL of sterile 3% thioglycolate solution (Cat. 225650, BD, USA) using a 1 mL syringe fitted with a 26G needle. Three to four days after the injection, the mice were euthanized, and 10 mL of sterile PBS was carefully injected into the peritoneal cavity using a syringe. The abdomen was gently massaged to dislodge adherent cells. The peritoneal lavage fluid was aspirated with a syringe and transferred to sterile centrifuge tubes. The lavage fluid was centrifuged at 300 × g for 5 minutes at 4°C to pellet the cells. The supernatant was discarded, and the cell pellet was resuspended in fresh sterile PBS or an appropriate cell culture medium.

### Phagocytosis Assays

RAW264.7 cells and primary macrophages were seeded at a density of 10⁵ cells per well in a 96-well plate containing DMEM-LG (5.5 mM glucose) or DMEM-HG (30 mM glucose) medium supplemented with 10% FBS. The cells were incubated at 37°C in a 5% CO₂ atmosphere for 24 hours. E. coli was then cocultured with the macrophages at a multiplicity of infection (MOI) of 100 for 3 hours in antibiotic-free DMEM-LG or DMEM-HG medium. After the coculture, the supernatant was collected for the phagocytosis assay. The supernatant was serially diluted in PBS to 10⁻⁴ and 10⁻⁵ and plated onto LB agar. The phagocytosis rate was calculated as the ratio of phagocytosed bacteria to the total bacteria.

### Statistical Analysis

Statistical analysis and graph generation were conducted using GraphPad Prism 9 (GraphPad Software, Inc., La Jolla, CA, USA) or R version 4.2.3. Results are presented as mean±SEM.

## RESULTS

### Cardiac Injury-elevated Gut Permeability Was Worsened by Hyperglycemia

To evaluate the impact of metabolic dysfunction on gut and heart interactions, we employed a hyperglycemia model induced by streptozotocin (STZ) injection and a high-fat, high-sucrose diet (HFHSD) and assessed gut permeability following MI using FITC-dextran assays (Figure 1A and 1B). Post-MI mice showed a marked increase in gut permeability, with hyperglycemia further amplifying this effect (Figure 1C). Alcian Blue staining revealed a significant reduction in the thickness of the intestinal mucus layer in post-MI mice (Figure 1D; Figure S1). The thickness of intestinal mucus was further reduced in hyperglycemic mice post-MI (Figure 1D; Figure S1). Further examination of the intestinal epithelium revealed a substantial downregulation of the tight junction protein Claudin-5 (Figure 1E) in post-MI mice. This reduction was also exacerbated in hyperglycemic conditions, highlighting the critical effect of hyperglycemia on the tight junction integrity in intestine. These results collectively underscore a synergistic impact of hyperglycemia and cardiac injury on gut barrier integrity.

**Figure 1.**
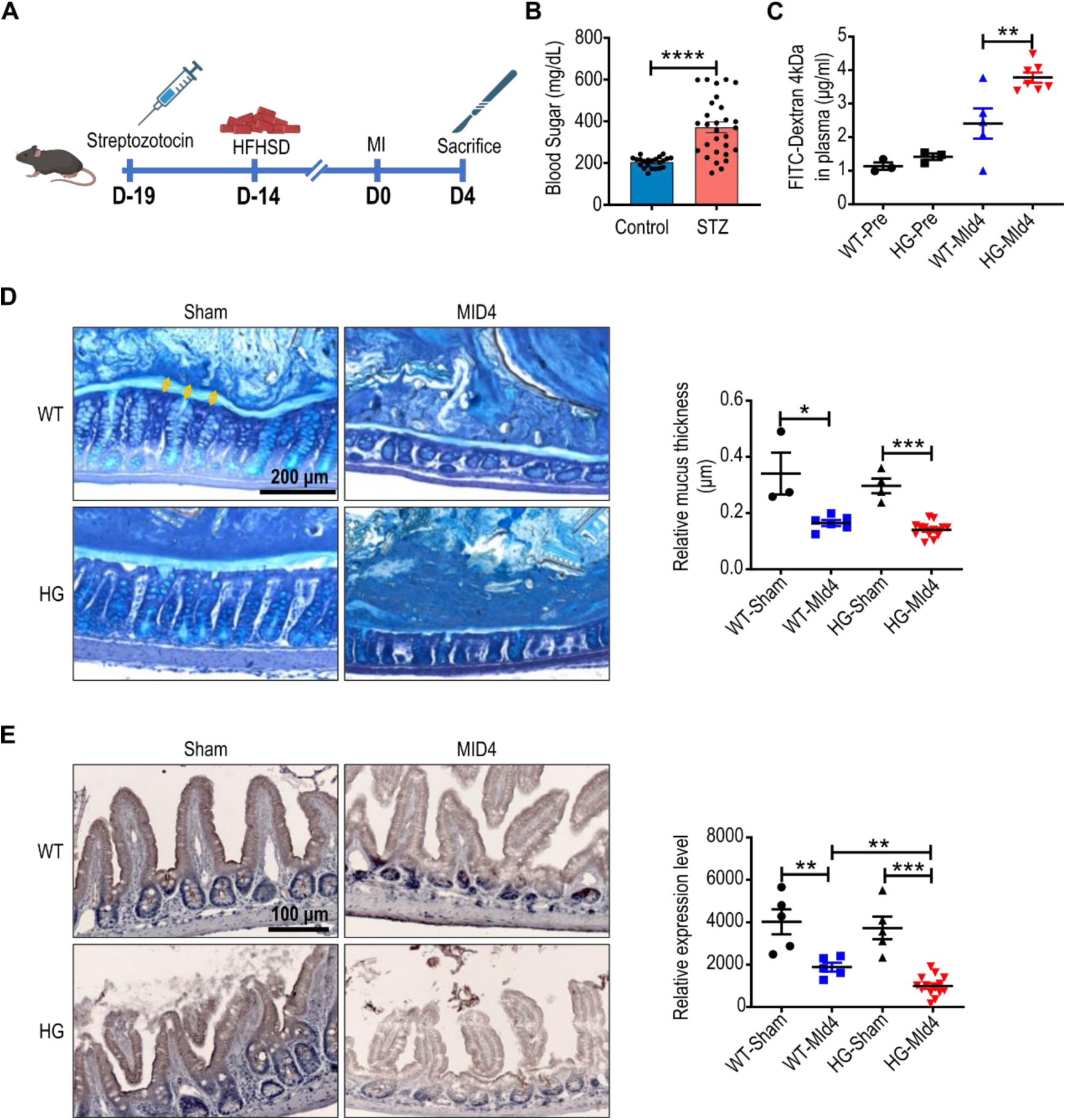
Cardiac injury alters gut permeability. **A.** Schematic representation of the experimental procedure for inducing hyperglycemia (HG) and cardiac injury. **B.** Quantification of blood glucose levels in mice following streptozotocin (STZ) injection and high-fat, high-sucrose diet (HFHSD) feeding. **C.** Quantitative analysis of FITC-dextran translocation in wild-type (WT) and HG mice after MI, indicating changes in gut permeability. **D.** Representative images of colon sections stained with Alcian Blue, along with quantification of colon mucus thickness in WT and HG mice post-MI. **E.** Representative immunohistochemistry images and quantification of the tight junction protein Claudin-5 in colon tissue. Data are presented as mean ± SD. Statistical analysis was performed using t test. *p < 0.05, **p < 0.001, ***p < 0.0001.

### Hyperglycemia Accelerated Gut Microbiota Translocation and Liver Colonization post-MI

To explore the dynamics of microbial translocation following MI and the impact of metabolic dysfunction, we analyzed organ samples collected at multiple time points post-MI. Anaerobic cultures from various organs showed substantial bacterial growth, with the liver emerging as the primary site of colonization (Figure 2). The bacterial load in the liver peaked on day 4 post-MI (Figure 2A and 2B). In hyperglycemic mice, bacterial translocation to the liver occurred earlier and was markedly enhanced, beginning on day 1 post-MI and persisting through day 10 (Figure 2A and 2C). Notably, hyperglycemia significantly accelerated the translocation of gut-derived microbiota, as evidenced by higher positive rate of bacterial culture in the liver and other organs compared to normoglycemic controls (Figure 2A and 2C). These results highlight the vital interplay between metabolic dysfunction and cardiac injury in promoting gut-derived bacterial dissemination, with hyperglycemia exacerbating the extent and persistence of microbial colonization in extraintestinal organs.

**Figure 2.**
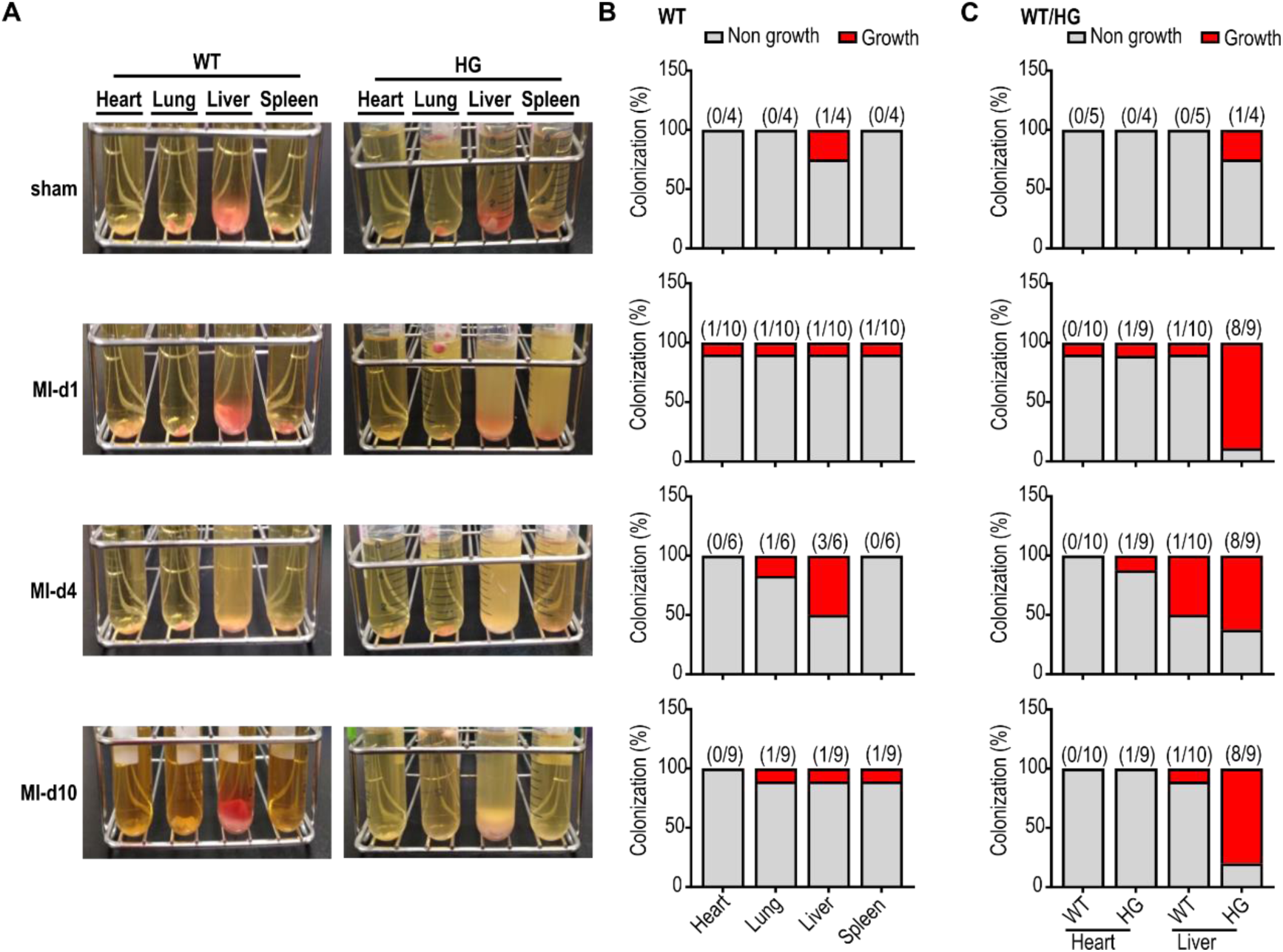
Hyperglycemia enhances the translation of gut microbiota into other organs. **A**. Representative images of tissue cultures from the heart, lungs, liver, and spleen grown in BHI broth under anaerobic conditions. The images illustrate bacterial growth in various organs, highlighting the effect of hyperglycemia on gut microbiota translocation. **B**. Quantification of bacterial culture positive rates in organs from WT mice post-MI. **C**. Quantification of bacterial culture positive rates in heart and liver from WT and HG mice post-MI. WT: wild type; HG: hyperglycemia.

### Gut-derived Bacterial Colonization in Cardiac Tissue

Bacterial dissemination was observed in multiple organs post-MI, with the liver being the primary site of colonization rather than the heart. To investigate whether bacteria could colonize the myocardium despite its continuous blood flow, microbiota cultured from stool samples were directly injected into the myocardium (Figure S2A). Organ sample culture confirmed robust bacterial colonization within the myocardium, reaching a critical threshold of 10⁴ CFU (Figure S2B and S2C). The data above directly demonstrated that gut-derived bacteria could establish colonies within cardiac tissue under suitable conditions.

### Inflammatory Cytokines Promote Microbial Dissemination following Cardiac Injury

High-throughput cytokine screening revealed significantly elevated levels of inflammatory cytokines, particularly TNF-α, following MI (Figure 3A). We then investigated whether TNF-α is competent in promoting microbial dissemination (Figure 3B). Treating mice with TNF-α in the context of hyperglycemia showed its ability to independently induce the translocation of gut-derived microbiota into extraintestinal organs (Figure 3C through 3E). However, the extent of dissemination triggered by TNF-α alone was less pronounced compared to that observed in the setting of MI-induced injury, which involves additional systemic factors and mechanisms (Figure 2C and 3E). These findings suggest the pivotal role of inflammatory cytokines, such as TNF-α, in mediating gut barrier dysfunction and microbial translocation.

**Figure 3.**
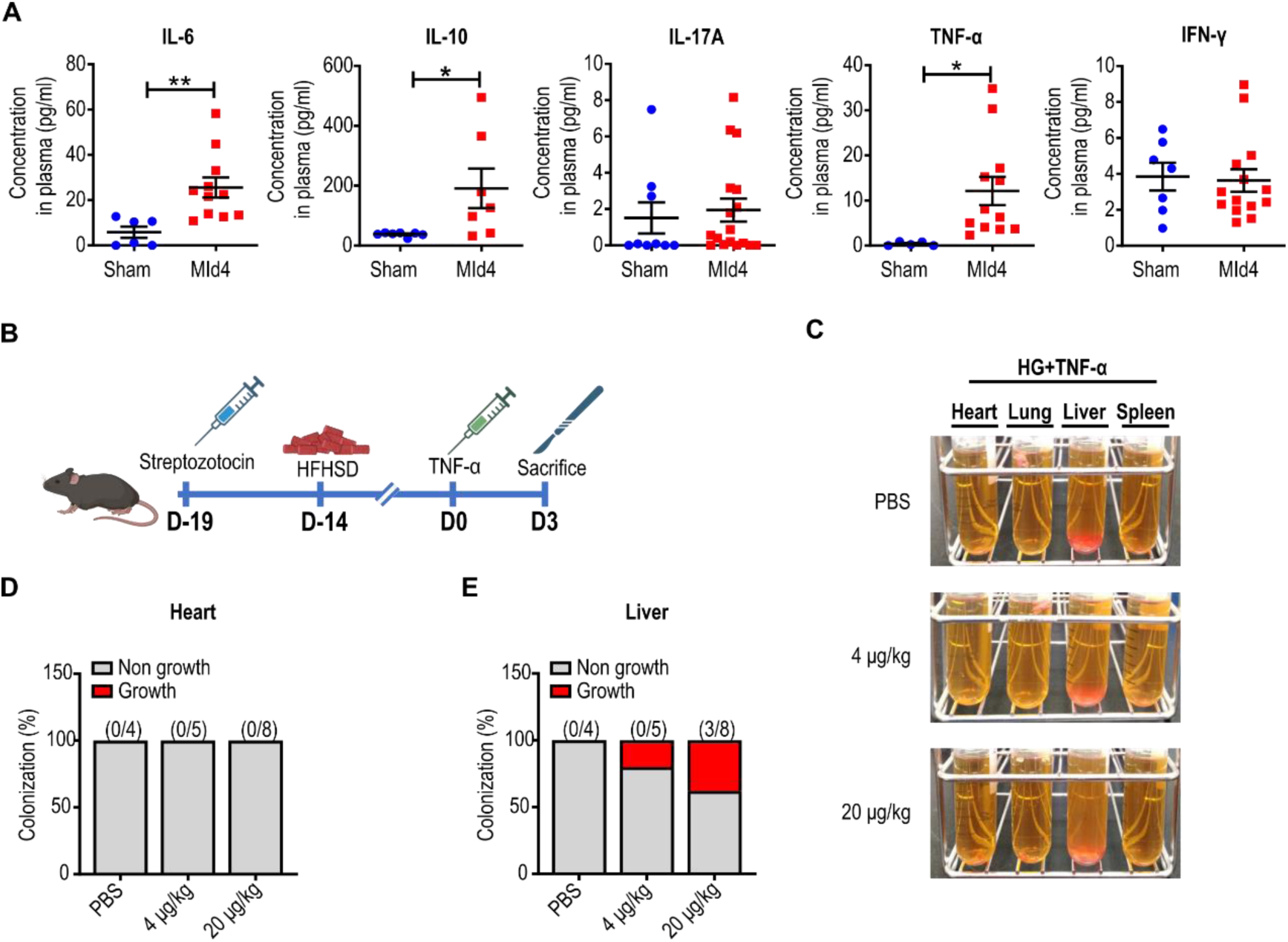
Cardiac injury stimulates the production of inflammatory factors. **A**. Cytokine levels measured in mouse plasma after MI using high-throughput screening. The data demonstrate the elevated production of inflammatory factors in response to cardiac injury. Data are presented as mean ± SEM. Statistical analysis was performed using unpaired t test. *p < 0.05, **p < 0.001, ***p < 0.0001. **B**. Schematic representation of the experimental procedure for TNFα-induced microbiota dissemination in hyperglycemia mice. **C**. Representative images of tissue cultures from the heart, lungs, liver, and spleen grown in BHI broth under anaerobic conditions. **D** and **E**. Quantification of bacterial culture positive rates in heart and liver from hyperglycemic mice following TNF-α injection. Tissue cultures from hyperglycemic mice treated with or without TNF-α injection.

### Hyperglycemia Impairs Hepatic Macrophage Function, Facilitating Microbial Dissemination and Interception in the Liver post-MI

The gut-liver axis, a bidirectional connection facilitated by the portal vein, plays a crucial role in regulating microbial translocation and immune responses. The liver, equipped with an extensive reticuloendothelial system, acts as a primary defense mechanism by using macrophages to engulf and eliminate foreign materials [33]. Using a FITC-labeled bacteria tracking system (intravital imaging), we observed that gut bacteria entering the bloodstream were intercepted by the liver through the portal vein (Figure 4; video 1). Bacterial presence in the portal vein blood peaked one day after MI and declined by day three (Figure 4B). However, in hyperglycemic mice, this interception was significantly impaired, leading to higher bacterial colonization in the liver. This deficiency was linked to reduced macrophage phagocytic activity.

**Figure 4.**
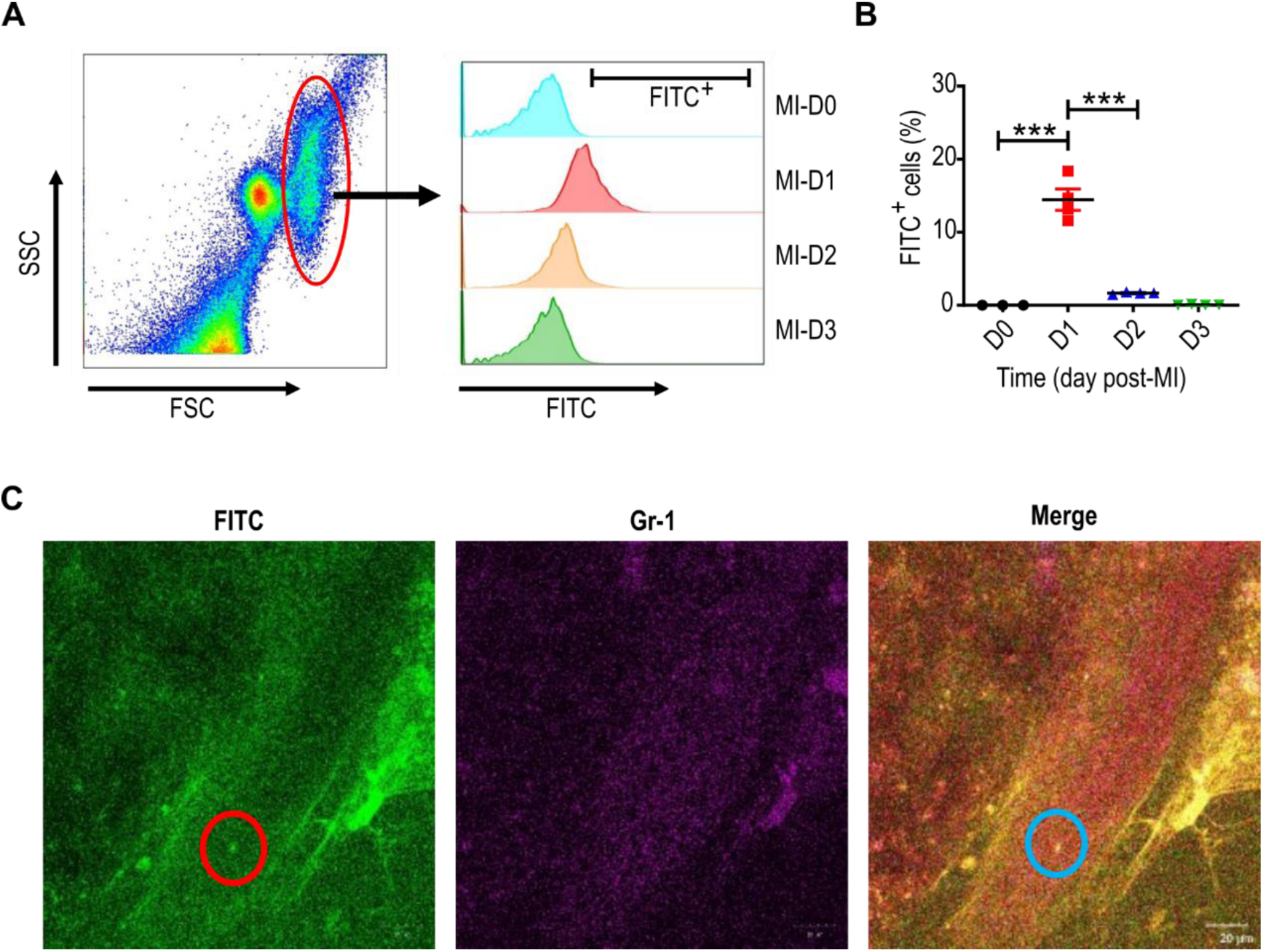
Disseminated gut microbiota translocate to the liver through the portal vein. **A.** Flow cytometry measurement of the percentage of FITC+ bacteria in blood collected from the portal vein after MI. **B.** Quantification of FITC+ bacteria percentage in portal vein blood, analyzed using flow cytometry post-MI. **C.** Representative intravital microscopy images showing FITC-labeled bacteria traveling through the portal vein. Data are presented as mean ± SD. Statistical analysis was performed using Kruskal-Wallis test, followed by Dunn’s Correction. ***p < 0.0001.

Hyperglycemia was found to activate the NLRP3 inflammasome signaling pathway, which diminished the phagocytic capacity of liver macrophages [35]. Real-time PCR analysis revealed increased NLRP3 expression in macrophages under high-glucose conditions (Figure S3A), while functional assays demonstrated reduced phagocytic activity both in vitro and ex vivo (Figure S3B). This dysfunction allowed more bacteria to accumulate in the liver and facilitated their spread to other organs. Experiments using clodronate liposomes to deplete immune cells confirmed the importance of macrophages in controlling microbial translocation (Figure 5A, 5B). When macrophages were absent, bacterial colonization in the liver decreased, but the bacteria were redirected to other organs, such as the lungs, at later stages (Figure 5C, 5E, and 5F).

**Figure 5.**
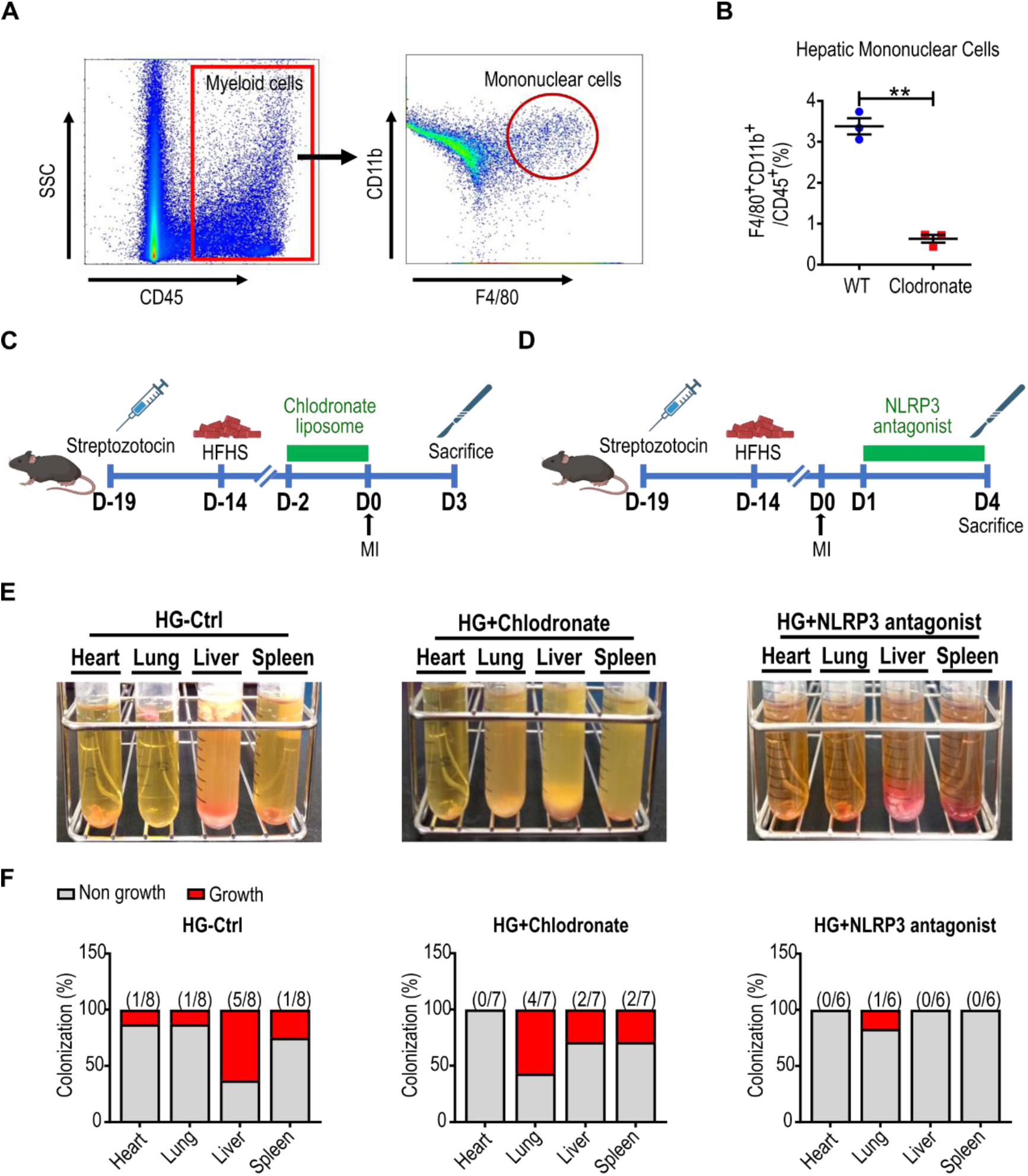
Hyperglycemia impairs the phagocytotic activity of macrophages. **A.** Flow cytometry analysis of the mononuclear cell population in hepatic cells. **B.** Quantification of hepatic mononuclear cells analyzed by flow cytometry in wild-type (WT) mice and mice injected with clodronate liposome. **C-D.** Schematic representation of the experimental procedure for chlodronate liposome (**C**) and NLRP3 antagonist (**D**) treatment in hyperglycemia mice. **E.** Representative images of tissue cultures from the organs grown in BHI broth under anaerobic conditions. Tissue cultures from hyperglycemic mice treated with or without chlodronate liposome or NLRP3 antagonist post-myocardial infarction. **F.** Quantification of bacterial culture positive rates in organs from hyperglycemic mice treated with or without chlodronate liposome or NLRP3 antagonist post-myocardial infarction. Data in (**B**) are presented as mean ± SEM. Statistical analysis was performed using unpaired t test. *p < 0.05, **p < 0.001, ***p < 0.0001.

Importantly, treatment with an NLRP3 antagonist restored macrophage phagocytic function, reducing bacterial buildup in the liver without increasing their spread to other organs (Figure 5D–5F). These findings highlight the vital role of liver immune function in preventing microbial dissemination, particularly in the context of MI and hyperglycemia. They underscore the need for targeted therapies to counteract hyperglycemia-induced immune dysfunction, which can have widespread systemic consequences.

### Stochastic Translocation of Gut Microbiota to Liver with Variable Genera Composition after Cardiac Injury

To determine whether the gut microbiota intercepted by the liver had a specific composition, two independent batches of liver- and stool-extracted samples were analyzed using 16S full-length sequencing. Remarkably, the dominant microbial genera in liver-extracted samples differed entirely between the two batches (Figure 6). Batch 1 was characterized by a higher prevalence of *Bacteroides acidifaciens* and *Limosilactobacillus reuteri*, whereas Batch 2 was dominated by *Acinetobacter junii* and *Comamonas denitrificans*. This variability indicates that while the liver consistently intercepts gut-derived microbiota post-MI, the specific genera that translocate and establish in the liver are likely influenced by stochastic, host-specific and environmental factors. These findings highlight the dynamic and variable nature of gut microbiota dissemination to the liver, suggesting that the process is largely driven by individual host conditions and external variables, ultimately shaping the microbial composition in the liver.

**Figure 6.**
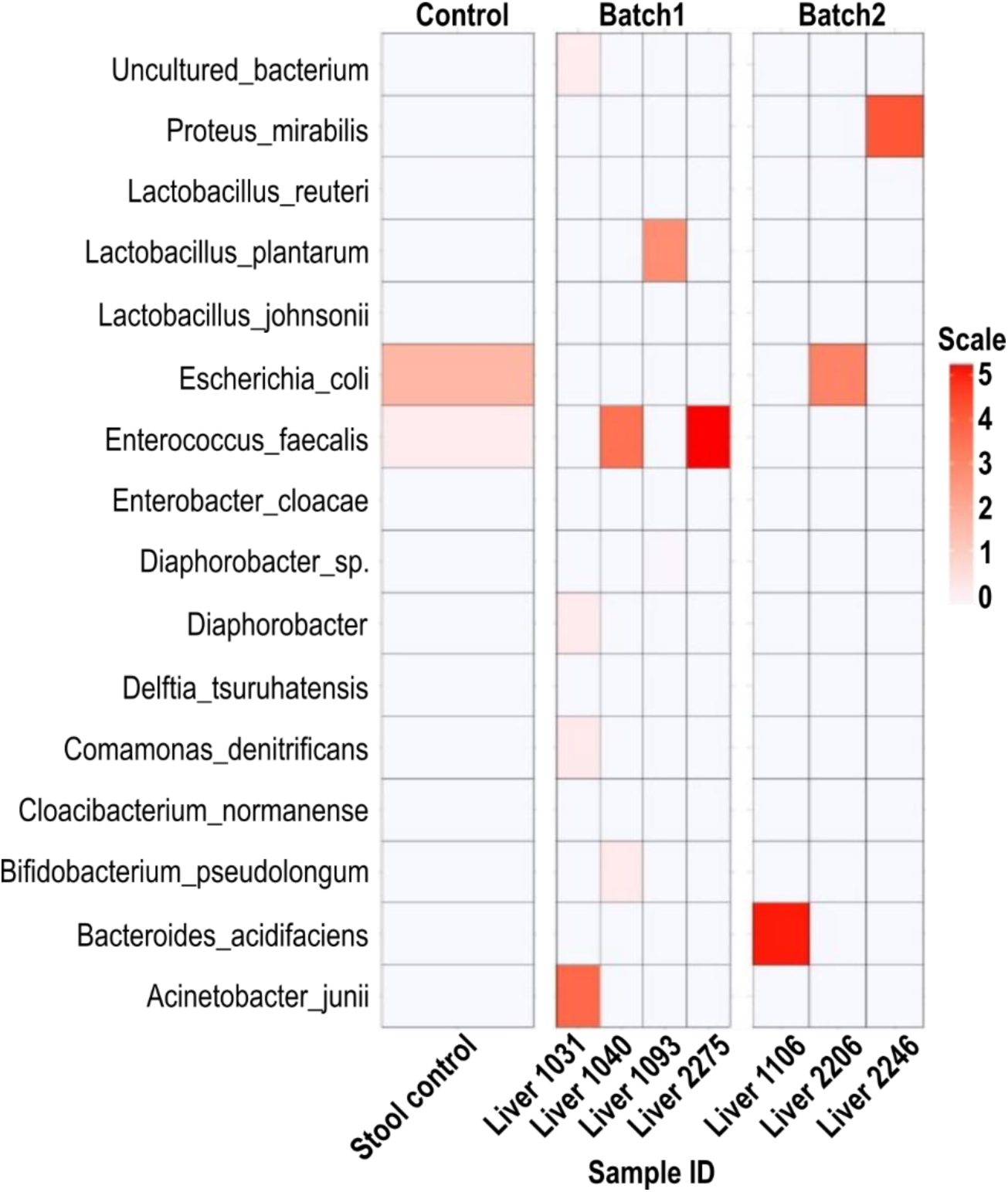
The dissemination of gut microbiota to the liver occurs randomly. Heatmap of the relative abundance of bacterial taxa identified across different liver samples and a stool control. The intensity of the color corresponds to the scaled abundance values, as indicated by the color gradient legend. The stool control serves as a reference for gut microbiota composition.

## DISCUSSION

Cardiac injury, such as MI, induces profound systemic effects that extend beyond the heart [9–11]. Among these, the disruption of gut barrier integrity has emerged as an essential factor in post-MI complications. Our study demonstrated a marked increase in gut permeability following MI, as evidenced by FITC-dextran assays and histological analyses (Figure 1). The observed thinning of the intestinal mucus layer and the reduced expression of the tight junction protein Claudin-5 underline the structural and functional compromise of the gut barrier. This phenomenon aligns with previous findings that MI-induced systemic inflammation and hypoxia disrupt the gut epithelium, facilitating the leakage of bacterial and inflammatory mediators into systemic circulation [18,19,32]. Hyperglycemia further exacerbates these effects, amplifying the degree of gut barrier dysfunction. The importance of glucose management after MI is well-documented [28,31], and this is particularly relevant to the observations in our study.

Effective glucose management after MI is critical, as research consistently links post-MI hyperglycemia—whether transient or persistent—to higher mortality and increased complications in both diabetic and non-diabetic patients [29,30]. Persistent hyperglycemia is associated with worse outcomes, while early intensive glucose control has been shown to improve survival by mitigating the systemic effects of hyperglycemia, including its detrimental impact on gut barrier integrity and immune regulation [28]. Hyperglycemia exacerbates oxidative stress and destabilizes tight junction proteins [32,33], leading to increased gut permeability and a heightened risk of systemic bacterial dissemination. These mechanisms underscore the need for a comprehensive approach to MI management that incorporates gut barrier-targeted therapies, such as probiotics, prebiotics, and agents that reduce intestinal permeability, particularly in patients with metabolic syndrome. By addressing both metabolic dysregulation and its downstream effects, this integrated strategy could significantly enhance recovery outcomes.

The liver, acting as an immune sentinel, intercepts gut-derived microbes that translocate via the portal vein. Our findings reveal that bacterial translocation to the liver peaks at day 4 post-MI, reflecting the liver’s indispensable role in mitigating systemic dissemination (Figure 4). However, hyperglycemia significantly accelerates this process, with earlier and more persistent bacterial colonization observed in hyperglycemic conditions, all the way through day 10 post-MI (Figure 2). This enhanced microbial translocation is likely driven by hyperglycemia-induced impairment of gut barrier integrity and hepatic immune defenses. This impairment compromises the liver’s ability to function as an effective immune barrier, thereby allowing greater systemic dissemination of gut-derived microbes. These findings highlight the need for metabolic regulation in post-MI care, with strategies aimed at reducing hyperglycemia and enhancing hepatic immune function.

Our study provides direct evidence of bacterial colonization within myocardial tissue through direct injection (Figure S2). While hypoxic and necrotic regions caused by injury create a microenvironment conducive to bacterial growth in the heart, as previously observed in other hypoxic tissues [24,25], our findings show that the percentage of colonization in the myocardium remains very low post-MI (Figure 2). This suggests that myocardial colonization is not a major pathway for bacterial presence following MI. Instead, the liver intercepts most, if not all, translocated bacteria, effectively preventing significant bacterial colonization in the myocardium. The limited colonization observed raises questions about the broader clinical relevance of bacterial presence in the myocardium post-MI. Nevertheless, even minimal bacterial infiltration could trigger localized inflammation and potentially impair cardiac repair in susceptible cases. These findings underscore the importance of further research to elucidate the conditions under which myocardial bacterial colonization occurs and its impact on cardiac recovery. Targeted antimicrobial strategies may still be valuable in preventing systemic bacterial dissemination, particularly in patients with compromised gut barrier function or metabolic dysfunction.

Elevated levels of systemic inflammatory cytokines, particularly TNF-α, were observed following MI (Figure 3A). TNF-α has been shown to independently drive gut barrier dysfunction and microbial translocation [19,32]. Our findings indicate that TNF-α acts as both a marker and a mediator of systemic inflammation in the post-MI setting. Although TNF-α alone cannot replicate the full extent of bacterial dissemination observed in MI-induced injury (Figure 3C through 3E), it significantly amplifies gut permeability and systemic inflammatory responses. Targeting inflammatory cytokines represents a promising therapeutic avenue. Pharmacological inhibitors of TNF-α and other pro-inflammatory cytokines have shown efficacy in reducing systemic inflammation and improving outcomes in preclinical models [19,35]. Future studies should evaluate the clinical utility of these interventions in MI patients, particularly those with coexisting metabolic disorders.

Previous research has shown that the liver’s role as a key immune barrier is heavily dependent on the functionality of its resident macrophages, particularly Kupffer cells, and are central to bacterial clearance [33,35]. Hyperglycemia-induced activation of NLRP3 inflammasome signaling emerged as a major factor impairing the phagocytic capacity of hepatic macrophages in our study. This impairment leads to reduced bacterial clearance, increased liver colonization, and enhanced systemic dissemination. Real-time PCR analyses confirmed elevated NLRP3 expression under hyperglycemic conditions (Figure S3), and pharmacological inhibition of NLRP3 restored macrophage function and reduced bacterial colonization (Figure 5). These findings highlight the therapeutic potential of targeting NLRP3 signaling in hyperglycemic patients with MI. By restoring macrophage functionality, NLRP3 inhibitors could mitigate systemic inflammation and reduce the risk of secondary complications. Additionally, dietary and lifestyle interventions aimed at improving glucose regulation may serve as complementary strategies to enhance hepatic immune defenses.

One thing noteworthy is the variability in the composition of gut-derived bacteria colonizing the liver. HiFi 16S rRNA sequencing revealed significant inter-individual differences in microbial genera, reflecting the stochastic nature of bacterial translocation (Figure 6). This variability is likely influenced by host-specific factors such as baseline gut microbiota composition, immune responses, and environmental exposures [39,40]. These findings underscore the importance of personalized medicine in managing post-MI complications. Incorporating microbiome profiling into clinical care could help predict the risk of microbial translocation and identify patients who may benefit from targeted interventions. Future research should explore the mechanisms driving this variability and develop strategies to modulate gut microbiota composition for improved clinical outcomes.

## Conclusion

Our findings provide a comprehensive understanding of the interplay between cardiac injury, metabolic dysfunction, and microbial translocation. MI-induced disruption of the gut barrier facilitates bacterial dissemination, with hyperglycemia exacerbating these effects by impairing hepatic immune defenses. The liver plays a central role as an immune sentinel, intercepting translocated microbes; however, its capacity is compromised under hyperglycemic conditions. The variability in microbial translocation highlights the need for personalized approaches to post-MI care. By targeting gut barrier integrity, inflammatory cytokines, and NLRP3 signaling, we identify promising therapeutic avenues for mitigating systemic inflammation and improving outcomes in patients with cardiovascular and metabolic diseases. These findings advance our understanding of the gut-liver-heart axis and pave the way for innovative strategies to address the challenges posed by this complex interplay.

## List of abbreviations

MI: myocardial infarction
SPF: specific pathogen-free
BHI: brain heart infusion
HFHSD: high-fat, high-sucrose diet
PBS: phosphate-buffered saline
CCS: circular consensus sequence
DADA2: divisive amplicon denoising algorithm 2
LEfSe: linear discriminant analysis effect size
HRP: horseradish peroxidase
HBSS: Hank’s balanced salt solution
MOI: multiplicity of infection
CFU: colony-forming units
NLRP3: NOD-like receptor family pyrin domain containing 3
LPS: lipopolysaccharides
STZ: streptozotocin
CVD: cardiovascular diseases

## Acknowledgements

The authors would like to thank the Core Facilities at the Institute of Biomedical Sciences, Academia Sinica, for the support of FACS, high-content imaging, and animal studies. We are also grateful to Dr. Peilin Chen’s lab at the Research Center for Applied Sciences, Academia Sinica, for their technical assistance with real-time intravital imaging, which was supported by funding for the imaging system (AS-IA-110-M04).

## Authors’ contributions

P.C.H.H. supervised the research and acquired fundings; T.W.H.T. and H.C.C. designed and performed experiments, carried out data analysis, image analysis and wrote the manuscript; S.N.N.P, S.C.R., P.J.L., E.W, C.W.K, acquired data and specimen; P.C.H.H., P.C., Y.C.H., Y.C.C. and T.W.H.T. reviewed and edited the manuscript.

## Availability of Data and Materials

Data and materials in this study will be available upon request to the corresponding author Dr. Patrick C.H. Hsieh.

## Disclosures

### Ethics approval and consent to participate

This study did not involve human participants, human data, or human tissue. Therefore, ethics approval and informed consent were not required.

### Consent for publication

Not applicable.

