## Supplemental figure 1~3 for "Hyperglycemia-Driven Hepatic Immune Dysfunction Facilitates Microbial Dissemination Post-Myocardial Infarction"

Supplemental material includes: **Figure S1-S3**

Correspondence to Patrick C.H. Hsieh, MD, PhD, FAHA

Institute of Biomedical Sciences, Academia Sinica

128, Academia Road, Section 2, Nankang District, Taipei 115, Taiwan.

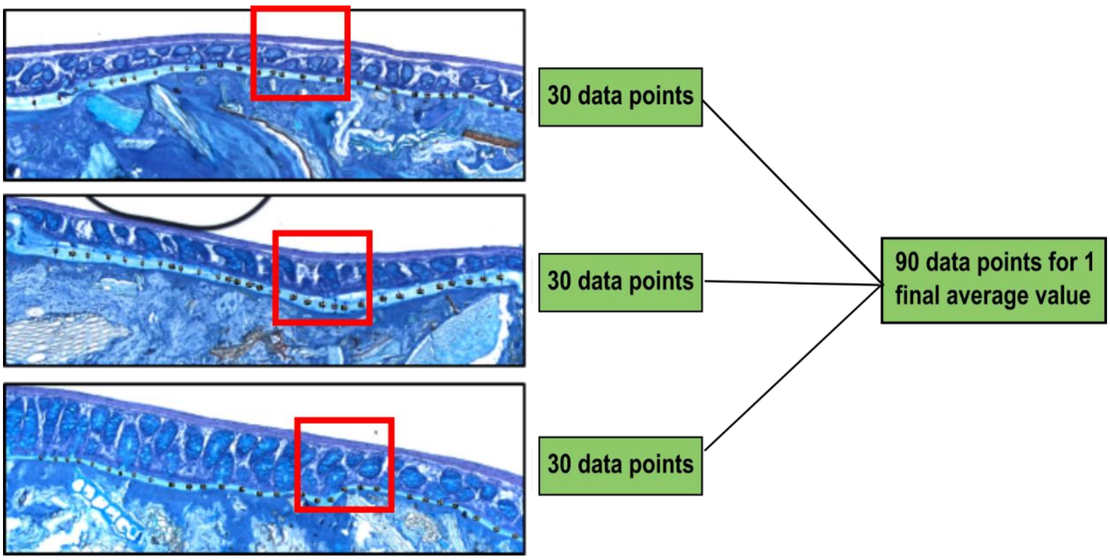


**Figure S1. Representative images showing the process of colon mucus thickness quantification.**

Images illustrate the methodology for measuring colon mucus thickness, including representative stained sections and the quantification procedure.

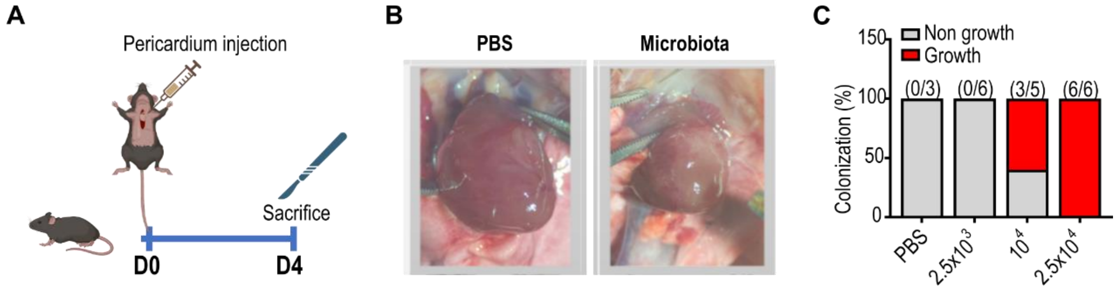


**Figure S2. Bacteria colonized in the heart after direct myocardial injection.**

**A.** Schematic representation of the experimental procedure used to examine bacterial colonization in the heart following direct bacterial injection. **B.** Representative in vivo heart images showing bacterial colonization after injection. **C.** Quantification of bacterial culture positive rates in heart samples from mice post-bacterial injection.

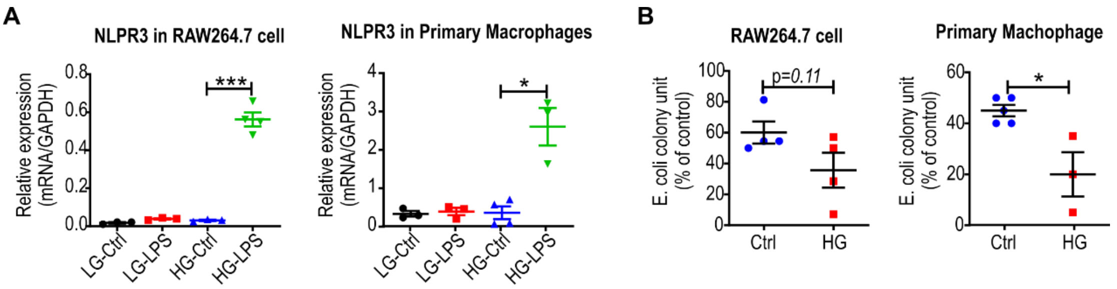


**Figure S3. Hyperglycemia-induced NLRP3 activation reduces phagocytic activity in macrophages**

**A.** NLRP3 RNA expression levels measured by real-time PCR in RAW264.7 cells and primary macrophages under lipopolysaccharide (LPS) stimulation. **B.** Assessment of glucose's impact on the phagocytic activity of RAW264.7 cells and primary macrophages as determined by CFU assays. Data are presented as mean ± SD. Statistical analysis was performed using t test. $*p < 0.05$, $**p < 0.001$, $***p < 0.0001$.
